# OMAnnotator: a novel approach to building an annotated consensus genome sequence

**DOI:** 10.1101/2024.12.04.626846

**Authors:** Sadé Bates, Christophe Dessimoz, Yannis Nevers

## Abstract

**Motivation:** Advances in sequencing technologies have enabled researchers to sequence whole genomes rapidly and cheaply. However, despite improvements in genome assembly, genome annotation (i.e. the identification of protein-coding genes) remains challenging, particularly for eukaryotic genomes: it requires combining several approaches (typically *ab initio*, transcriptomics, and homology search), each with its own pros and cons. Deciding which gene models to retain in a consensus is far from trivial, and automated approaches tend to lag behind laborious manual curation efforts in accuracy.

**Results:** Here, we present OMAnnotator, a novel approach to building a consensus annotation. OMAnnotator repurposes the OMA algorithm, originally designed to elucidate evolutionary relationships among genes across species, to integrate predictions from different annotation sources, using evolutionary information as a tie-breaker. We validated OMAnnotator by reannotating the *Drosophila melanogaster* reference genome and comparing it with the expert-curated reference and results from the automated pipelines BRAKER2 and EvidenceModeller. OMAnnotator produced a consensus annotation that outperformed each individual input and surpassed the existing pipelines. Finally, when applied to three recently published genomes, OMAnnotator gave substantial improvements in two cases, and mixed results in the third, which had already benefited from extensive expert curation.

**Conclusion:** We introduce an original, flexible, and effective approach to annotating genomes by integrating multiple lines of evidence. The method’s robustness is underlined by its successful implementation in re-annotating recently published genomes, opening up new avenues in eukaryotic genome annotation.

## 1. Introduction

With the advances in sequencing technology, sequencing a genome is faster and more affordable than ever. However, annotating the increasing number of newly sequenced genomes remains labour-intensive and error-prone (Salzberg 2019; Meyer *et al*. 2020; Scalzitti *et al*. 2020). Genome annotation involves identifying the features in a genome sequence, including protein coding or non-coding genes, regulatory elements and structural variants such as inversions and repeats, which is essential for understanding its underlying biology. Here, we focus on the prediction of protein coding genes and describe a new approach to improve the ease and accuracy of this process.

Three major classes of methods are used to predict genes in genome assemblies: *ab initio* gene prediction, transcript alignment and homology alignment (Hoff and Stanke 2015; Mudge and Harrow 2016). The first of these involves using a gene finder algorithm, such as AUGUSTUS (Stanke *et al*. 2008), to identify genes in a genome assembly based on gene structures in a model species it has been trained on. Transcript alignment is the alignment of reads from RNA sequencing to the reference assembly to localise transcripted regions of the genomes. Finally, homology alignment involves searching for regions in the newly sequenced genomes that are similar to coding genes in a closely related species to identify likely homologous genes.

One of the main challenges in genome annotation is combining the genes predicted by different—and sometimes contradictory—annotation methods into a consensus. A good consensus annotation accurately captures most genes in the genome, i.e., it retains the maximum number of true gene predictions while dropping false gene predictions. Genome annotation pipelines such as BRAKER2 (Brůna *et al*. 2021) aim to achieve this using RNAseq or protein sequence evidence during the AUGUSTUS (Stanke *et al*. 2008) iterative training and gene prediction processes, which improves annotation accuracy compared to using AUGUSTUS alone. Nevertheless, combining gene models from multiple evidence sources tends to be needed to remove false predictions from the consensus annotation. Existing approaches such as EVidence Modeller (EVM) (Haas *et al*. 2008) produce a consensus annotation by assigning quality weightings to annotation sets produced by different annotation methods. Although this increases the specificity of the annotation by dropping false predictions from a lower-quality set, less abundant true gene predictions may also be excluded from the consensus.

To approach this long-standing issue from a new angle, we sought to build consensus gene sets using an approach developed to model genome evolution. Our tool ‘OMAnnotator’ repurposes OMA (Orthologous MAtrix) standalone (Altenhoff et al., 2019), a state-of-the-art orthology inference software, to reconstruct a consensus annotation set from an arbitrary number of input annotation sets. OMA standalone infers Hierarchical Orthologous Groups (HOGs), sets of genes within a taxonomic range that descended from the same gene in the common ancestor of the taxon (Train *et al*. 2017; Zahn-Zabal, Dessimoz and Glover 2020). OMAnnotator combines annotations from different sources through its HOG inference software as if they were closely related species, with the consensus annotation being equivalent to the common ancestor of the annotation sets. By doing this, it can select gene models supported by multiple lines of evidence: agreement between predictions from the user-provided annotation sets (related by sequence similarity) and the existing annotations of related species (Figure 1). All genes retained in the consensus are supported by at least two lines of evidence (orthology with another annotation method and/or other species).

**Figure 1:**
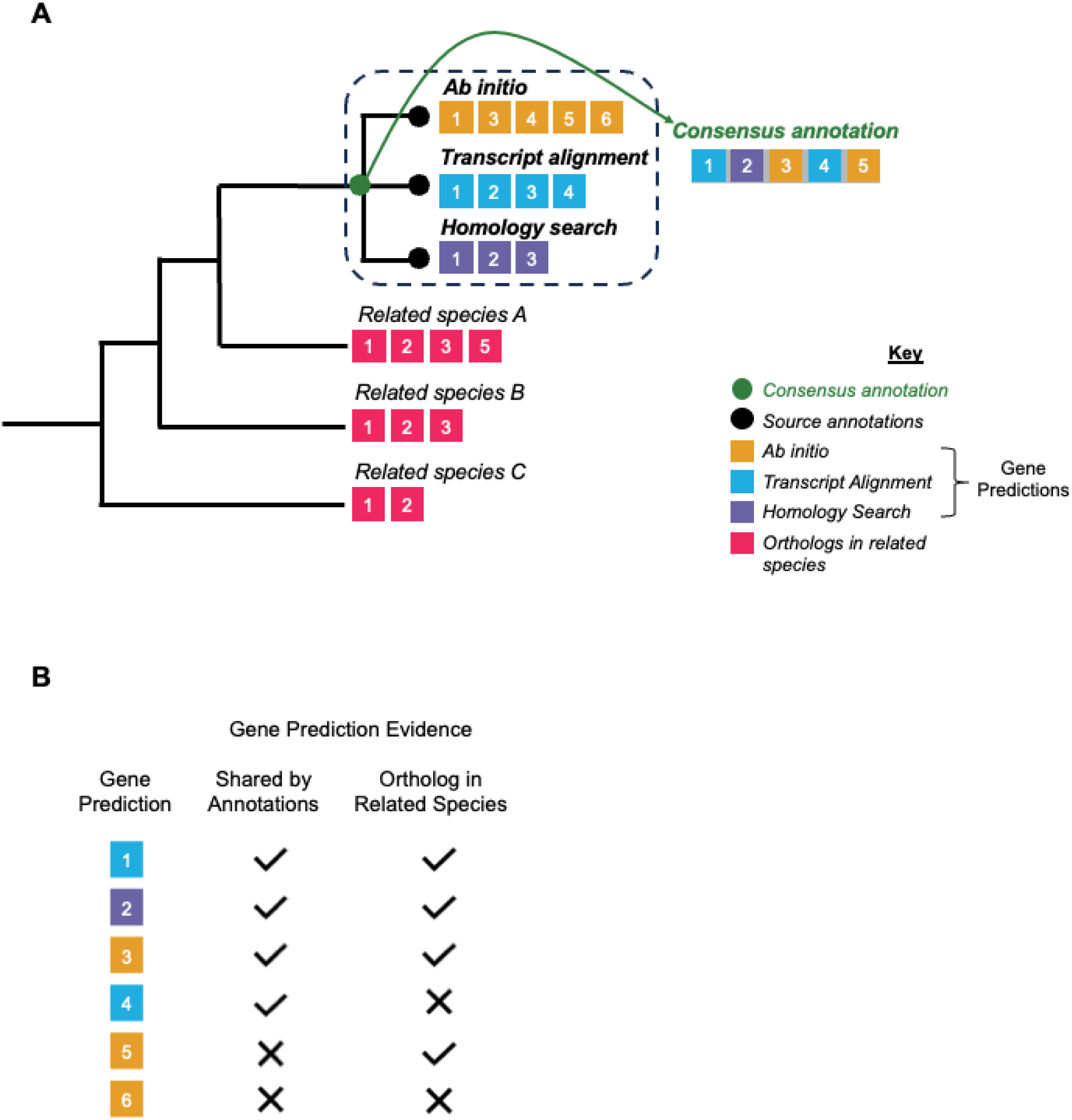
Overview of the OMAnnotator approach. **A**) An example of a simple species tree inputted to OMA is shown on the left. The numbered yellow, blue, and purple squares represent gene predictions from source annotations, and the numbered red squares represent their orthologs in related species. Species trees should comprise ∼10 species (downloaded from the OMAbrowser) that are related to the target species (3 are shown here, for simplicity) and source annotations added as a polytomy. OMAnnotator combines predictions from individual source annotations (black circles) into a consensus (green circle), leveraging orthology evidence from related species where gene predictions are not shared between annotations. **B)** Summary of evidence for retaining gene predictions. The colour of the gene prediction reflects its source annotation. OMAnnotator retains gene predictions (numbered coloured squares) that are shared by the annotation sets (predictions 1-4), or have an ortholog in a related species (predictions 1-3 and 5). Where gene predictions are shared between annotations, the predictions with the longest sequence are retained.

We validated this approach by using it to re-annotate the latest *Drosophila melanogaster* genome assembly, starting from the ground up. OMAnnotator integrated three lines of evidence: de novo gene prediction, RNA-seq data, and homology search. OMAnnotator’s output was then compared to the high-quality reference annotation available for this model organism. We also compared the OMAnnotator results to those obtained using EVM and BRAKER2 on the same input data. Finally, we used OMAnnotator to re-annotate the genomes of three species previously annotated with EVM.

## 2. Methods

### 2.1. Description of the OMAnnotator pipeline

The OMAnnotator pipeline relies on the inference of orthologous groups provided by the OMA Standalone software (Altenhoff *et al*., 2019) to combine gene predictions from different annotation methods into a consensus annotation. The pipeline takes GFF3 files produced by any annotation methods as its primary input, uses them to form a consensus annotation and outputs a combined annotation in GFF3 and FASTA format. It is executed in three main steps: setting up annotation data, HOG inference and consensus annotation extraction (see Figure 2 for an overview).

**Figure 2:**
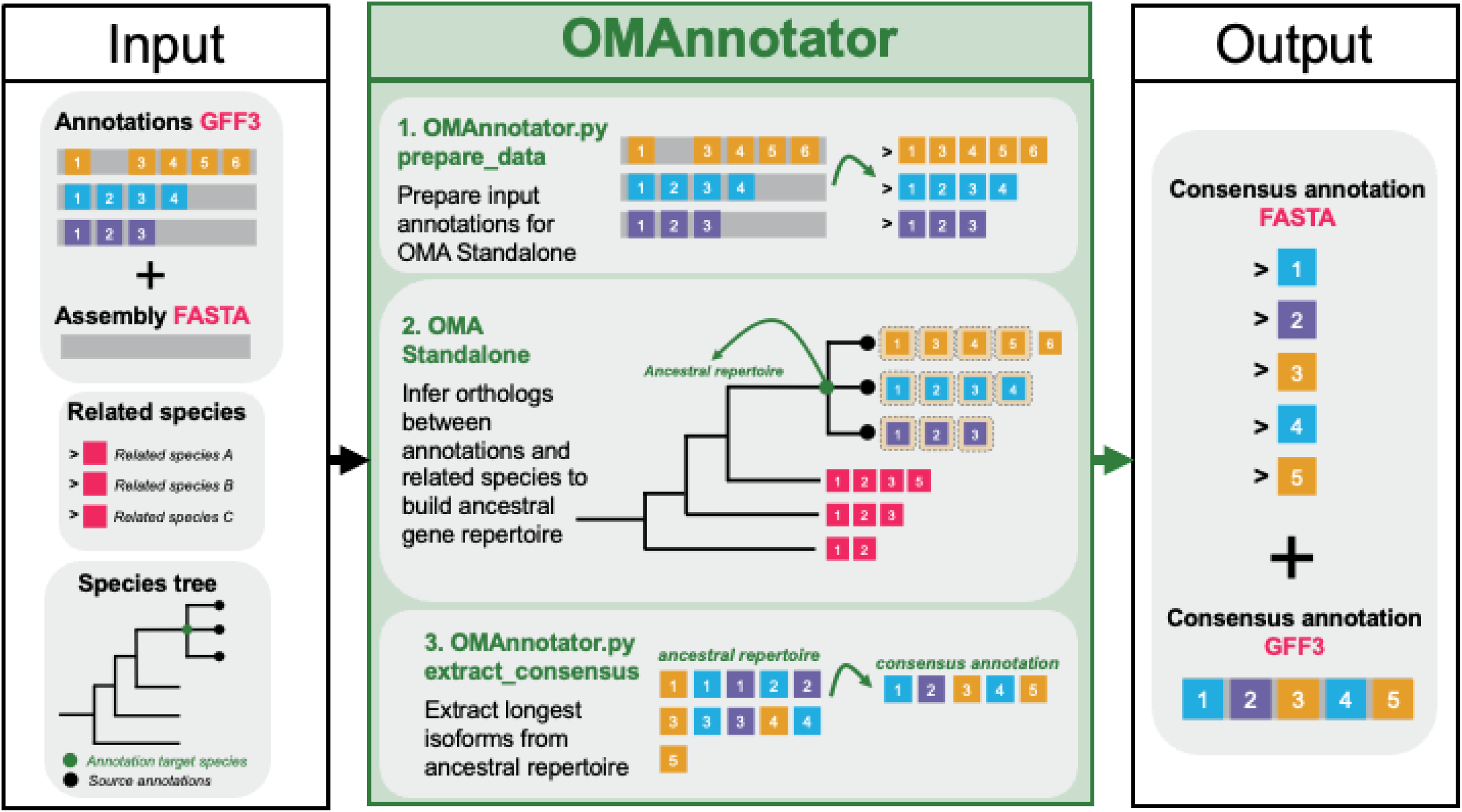
an overview of the OMAnnotator workflow. OMAnnotator takes as input: GFF3 format annotations from different sources, a FASTA format genome assembly, and a file of precomputed orthology relationships between a set of user-selected species that are related to the annotation target. This ‘related species’ file is downloaded from the OMA browser. **1)** The ‘prepare_data’ command from OMAnnotator.py prepares the input data for OMA Standalone. **2)** The user-specified species tree is added to the OMA Standalone parameters file. OMA Standalone infers orthology between the annotations (black circles) and related species at the ancestral node (green circle). **3)** Gene predictions supported by multiple annotations or that have an ortholog in a related species are extracted using the ‘extract_consenus’ command from OMAnnotator.py. Where gene predictions are shared between annotations, the prediction with the longest CDS is selected as representative in the consensus annotation. OMAnnotator outputs the consensus annotation in FASTA and GFF3 formats.

#### 2.1.1. Setting up annotation data for OMAnnotator

A local copy of the OMA Standalone software and a file of precomputed orthology relationships between a set of species are prerequisites. We recommend the species be selected to maximise taxonomic diversity and include species that are closely related to the species being annotated. Such a file can be downloaded from the ‘Download>Export All-All’ section of the OMA Browser (https://omabrowser.org/oma/export/), which downloads an archive of precomputed pairwise sequence comparisons between the proteins of the selected species. Both the OMA Standalone software and the precomputed orthology relationships are locally stored in what will be hereafter referred to as the ‘OMA folder’.

The first step of the pipeline involves extracting the information needed to run the OMA algorithm from the input GFF3 files and the target species’ genomic sequence (Figure 2). This process is automated through the “prepare_data” command of the OMAnnotator software, which takes as input a folder containing any number of GFF3 annotation files and the genome assembly sequence to which they correspond. As feature naming by different annotation software often deviates from the GFF3 specifications (Lincoln Stein 2020), we have added an optional ‘feature_type’ argument to our prepare_data function. This argument takes a tab-separated value file indicating which features in the annotation file correspond to a gene, transcript or CDS according to the GFF3 specifications, giving OMAnnotator the flexibility to process annotations from a variety of software without the need for the user to reformat their source annotations. The prepare_data command extracts all protein sequences described in a GFF3 file and adds its resulting FASTA file to the ‘DB’ (database) subfolder in the user’s OMA folder. If any gene is predicted to have multiple isoforms, alternative splicing information is added to the DB subfolder as a ‘Splice file’. This enables OMA Standalone to select the isoform sequence most similar to its detected homologs as the main representative of each gene. The prepare_data command also writes a species tree file for the user-selected related species to the parameters file in the OMA folder.

#### 2.1.2. OMA Standalone

Next, the OMA Standalone pipeline is run as described in Altenhoff *et al*. (2019, 2021) to infer orthology relationships between proteomes. For this purpose, the parameter file is edited to specify a species tree (in Newick format) that includes the various input annotation sets to be combined, and the user-selected related species. The leaves of the tree must share the same names as the annotation FASTA files in the DB folder. We recommend that users edit the Newick format string that the prepare_data command writes to the parameter file during the earlier prepare_data step.

#### 2.1.3. Consensus extraction

The last step of the pipeline is the extraction of a consensus annotation from the OMA Standalone. This is done using the “extract_consensus” command of the OMAnnotator software, which takes as input the species tree specified above and the HierarchicalGroups.orthoxml file outputted by OMA Standalone. It generates a protein FASTA file and a GFF file corresponding to the consensus annotation.

The software retains as consensus genes any gene that is present in the OMA inferred ‘ancestral genome’ of the different annotation methods. As such, the ancestral genome contains any sequence shared by at least two annotation methods or those predicted in one of the annotations as long as they have detected orthologs in any outgroup species. This allows combining genes inferred by multiple methods but disregarding the ones with low support. When multiple annotation methods predict a gene, the version with the longest coding sequence is selected as the representative sequence for the consensus set.

### 2.2. Proof of principle: Annotating the *Drosophila melanogaster* genome sequence with OMAnnotator

To validate the OMAnnotator approach, we used it to annotate the *D. melanogaster* genome sequence from scratch. We downloaded the latest genome assembly (genomic release 6.32) without annotations from FlyBase and used three annotation methods to predict genes. We then used OMA standalone to combine the resulting annotation sets into a single FASTA and GFF3 sequence file (Lincoln Stein 2020).

#### 2.2.1 Gene prediction using AUGUSTUS

We ran AUGUSTUS with the unannotated *D. melanogaster* genome as input and the “--species=fly” option to specify *D. melanogaster* gene model parameters. No output format flag was set, producing an annotation in AUGUSTUS’ native GTF format.

#### 2.3.2 Transcript alignment using StringTie

17-day *D. melanogaster* adult tissue RNAseq data generated with paired-end Illumina sequencing during a study of the *D. melanogaster* developmental transcriptome (Bryce Daines; 2010) were downloaded from NCBI SRA (accession number SRS065821). Sequences were joined in FASTQ format using the fastq-dump tool from the SRA toolkit v2.10.9, and their quality was checked using FastQC (*Babraham Bioinformatics - FastQC A Quality Control tool for High Throughput Sequence Data*, no date). The adapter sequences were then trimmed using Trimmomatic (Bolger, Lohse and Usadel, 2014) in Paired End mode with the parameters “-phred33 -threads 24 ILLUMINACLIP:RNAseq_data/TruSeq2-PE.fa:2:30:10 LEADING:3 TRAILING:3 SLIDINGWINDOW:4:15 MINLEN:36”. Next, reads were aligned to the reference genome using StringTie, producing a genome alignment file (Pertea *et al*. 2015). The longest open reading frames were identified using TransDecoder, producing a GFF3 file of likely peptide sequences.

#### 2.3.3 Homology search: using GeMoMa

*Anopheles gambiae* proteome data from the assembly AgamP4 (Sharakhova *et al*. 2007) was downloaded from UniProt on 25/10/2021. GeMoMa (Keilwagen, Hartung and Grau 2019) was used to infer *D. melanogaster* gene models based on the *A. gambiae* protein sequences, with the following options specified “-Xmx50g GeMoMaPipeline GeMoMa.Score=ReAlign AnnotationFinalizer.r=NO o=true”.

#### 2.3.4 OMAnnotator: using OMA Standalone orthology inference to form a consensus annotation

A file of precomputed orthology relationships between 25 species (listed in Supplementary Table S1) was downloaded from OMA Browser as described above. FASTA format and any accompanying splice files were produced using the prepare_data command of the OMAnnotator software as described above. A species tree of the 25 outgroup species and the three annotations clustered on a single branch within the *Drosophila* clade was specified as a Newick format species tree (see Figure S1) in OMA Standalone’s parameters file. OMA Standalone was then run as described above, and a consensus annotation set was produced using the extract_consensus command of the OMAnnotator software.

### 2.4. Analysing the quality of annotations

All annotations were compared to the *D. melanogaster* reference annotation genomic release 6.32 downloaded from FlyBase (Gramates *et al*. 2022). First, a comparison of gene counts between the annotations produced by each method and reference annotation was used as a first estimate of the specificity and sensitivity of each method. The completeness of the gene set produced by each method was then analysed with BUSCO (v5.4.2) (Manni *et al*., 2021) using the *Dipteran* gene set from the odb10 release. Finally, the quality of the gene structure annotations in each annotation set was also compared to the *D. melanogaster* reference annotation using ParsEval (Standage and Brendel, 2012).

Briefly, ParsEval compares annotations by pairwise alignments. It defines “gene loci”, the smallest genomic regions that capture all the annotations that overlap, compares them and computes summary statistics. The numbers of gene loci shared, unique to the reference and unique to the prediction were used to evaluate the sensitivity and specificity of each method. Comparing the number of genes per gene locus gave a further indication of closeness to the reference. The number of matching CDS structures indicated the quality of the exon-intron structural predictions by each method. As the original *D. melanogaster* reference annotation was incompatible with ParsEval due to the presence of a trans-spliced gene, the entries relative to this single gene were removed from the GFF3 annotation before comparison.

### 2.5. Comparing OMAnnotator with other annotation pipelines

To compare the performance of OMAnnotator with other consensus approaches, we compared its output with that of EVM and BRAKER2 (Haas *et al*. 2008; Brůna *et al*. 2021) when annotating the same *D. melanogaster* genome sequence with the same input annotations.

We reformatted the AUGUSTUS and homology annotations into GFF3 format required by EVM, using its in-built scripts ‘augustus_GFF3_to_EVM_GFF3’ and ‘GeMoMa_gff_to_gff3’. The RNAseq annotation did not require reformatting. We assigned each annotation a weight of ‘1’, as we had no evidence to suggest the quality of gene structures predicted varied consistently across annotation sources. The software was run with the parameters –segmentSize <100000> and –overlapSize <15000>. The quality of the resulting consensus annotation was assessed as described above.

We then performed gene annotation with BRAKER2, which has various modes for training gene predictors on different annotation inputs. We first used RNAseq annotation as input. Then, we reran the software adding the homology annotation to test whether including homology data as evidence for gene finder training improved BRAKER2 sensitivity. We also ran the software with the homology data only. Before performing quality assessments, we converted the BRAKER2 GTF output into GFF3 format using the ‘gtf_to_gff3.pl’ script from the GenomeTools package (Gremme, Steinbiss and Kurtz 2013), as its GFF3 files did not conform to GFF3 standards (Lincoln Stein 2020). Where necessary, redundant sequences were removed using Awk. Prior to BUSCO analyses, custom scripts were used to convert the correct GFF3 files into FASTA format and select the longest isoform per gene.

### 2.6. Using OMAnnotator to re-annotate other genomes

To evaluate the flexibility and robustness of OMAnnotator on other genomes, we used it to re-annotate three non-model organism genomes for which the intermediate annotation data had been made available: *Siraitia grosvenorii* (Monk Fruit) (Xia *et al*. 2018), *Phyllostomus discolor* (Pale Spear-Nosed Bat) (Jebb *et al*. 2020), and *Olea europaea* (European Olive Tree) (Cruz et al. 2016). The genomes of each of these organisms were originally annotated using EVM to combine source annotation sets. The authors of the original annotations kindly provided their input source annotations and the raw consensus output from EVM where available, else they provided their curated final consensus annotation.

For each re-annotation, ten related species with good-quality proteomes were selected to maximise coverage of orders around the target species (listed in Supplementary Table S1). OMAnnotator was then run with the GFF3 source annotations and genome assembly as input, with the species tree specified in the OMA Standalone parameter file (Supplementary Figures S3-S5). Finally, the OMAnnotator consensus annotations were compared with the authors’ sequences using gene counts and BUSCO analyses. As these species are non-model organisms, BUSCO scores from each annotation method were compared with the genome assembly BUSCO scores instead of a reference annotation.

## 3. Results

### 3.1. OMAnnotator produces a high-quality annotation

As a proof of principle, we used OMAnnotator to annotate the *D. melanogaster* reference genome assembly (FlyBase, Genomic Release 6.32) from scratch, with three source annotations generated using three primary annotation approaches: gene prediction with AUGUSTUS (Stanke *et al*. 2008), RNAseq transcript alignment with StringTie (Pertea *et al*. 2015), and homology search with GeMoMa (Keilwagen, Hartung and Grau 2019). We then compared all annotations to the *D. melanogaster* reference annotation (genomic release 6.32), which as one of the best annotated model organism genomes, serves as a gold standard. This *D. melanogaster* reference gene set contained 13,986 protein coding genes and 99.9% complete single-copy *Dipertan* BUSCO genes, implying a high level of completeness, with a minimal fraction (0.2%) of duplicated, fragmented or missing genes (Figure 3).

**Figure 3:**
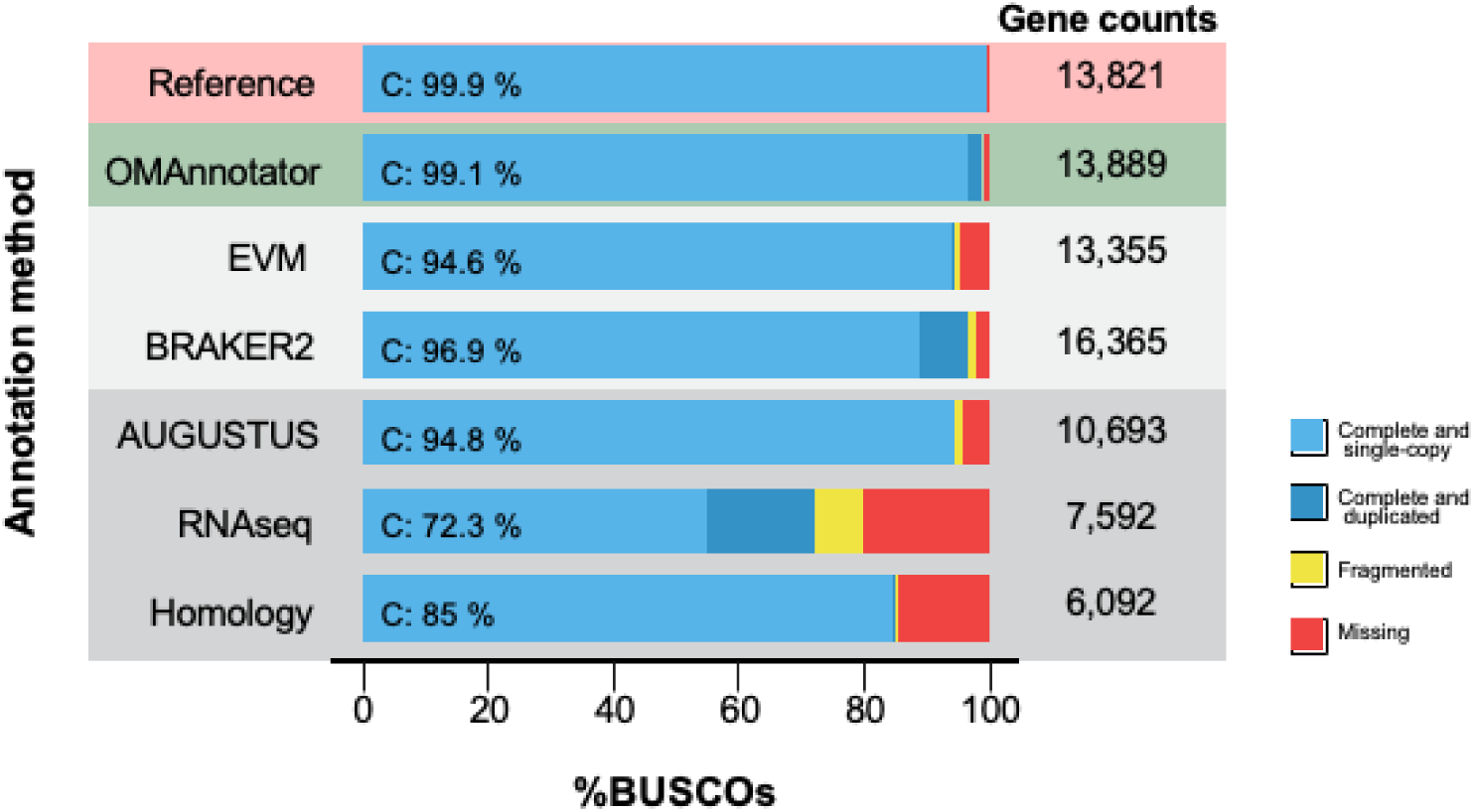
Results from BUSCO analyses of each annotation set are shown, along with the number of protein coding genes. Each annotation is compared to the gold standard reference annotation (FlyBase genomic release 6, version 4), highlighted in red in the first row. Individual annotations are highlighted in grey, and the consensus produced by OMAnnotator is highlighted in green. Consensus annotations produced by EVM and BRAKER2 are highlighted in light grey.

Of the individual methods, the RNAseq and homology annotations predicted the fewest genes (7,592 and 6,092 genes, respectively) in the assembly. Accordingly, they had more missing BUSCOs (RNAseq = 20.2% and homology = 14.3%; Figure 3). This was reinforced by ParsEval analysis, which found they had the highest percentage of gene loci (regions encompassing overlapping genes between the reference and prediction annotations) unique to the reference sequence (False Negative, FN) (RNAseq = 31.4% FN and homology = 38.4% FN; Table 1). In contrast, AUGUSTUS had the highest gene count of the individual methods, with 13530 genes – close to the reference gene count – but still had a notable proportion of fragmented and missing BUSCO genes (1.0% fragmented, 4.2% missing). It contained the fewest FNs of the individual annotation but also contained the most False Positives (FPs) at 13.9%.

**Table 1:** ParsEval results from pairwise comparisons between the D. melanogaster reference annotation (‘reference’) and the annotations predicted by each method are shown (‘prediction’). The ‘ParsEval comparison level’ column refers to the level at which the ParsEval features are being compared. Within this column, the ‘Gene loci’ are determined by ParsEval as the minimal genomic regions containing all overlapping annotations and the ‘CDS structure’ level is the coding sequence exon-intron structure of each annotation. The ‘Comparison’ column indicates how the ParsEval feature comparison was computed.

| *ParsEval*<br>*comparison level* | *Comparison* | *OMAnnotator* | *EVM* | *BRAKER2* | *AUGUSTUS*<br>*ab initio* | *StringTie*<br>*RNAseq* | *GeMoMa*<br>*Homology* |
| --- | --- | --- | --- | --- | --- | --- | --- |
| *Genes* | *Total number* | 13,889 | 13,077 | 16,365 | 10,693 | 7,592 | 7,575 |
| *Gene loci* | *Shared* | 86.64% | 83.26% | 79.10% | 83.21% | 68.40% | 61.31% |
|  | *Unique to*<br>*reference (FN)* | 10.59% | 4.27% | 4.03% | 2.92% | 31.37% | 39.19% |
|  | *Unique to*<br>*prediction (FP)* | 2.76% | 12.48% | 16.87% | 13.86% | 0.23% | 1.90% |
|  | *Genes per locus*<br>*reference* | 1.458 | 1.314 | 1.151 | 1.419 | 1.344 | 1.375 |
|  | *Genes per locus*<br>*prediction* | 1.448 | 1.251 | 1.425 | 1.365 | 0.982 | 0.756 |
| *CDS structure* | *Prediction CDS*<br>*matches to*<br>*reference* | 79.3% | 80.1% | 81.5% | 80.3% | 80.9% | 58.8% |

The OMAnnotator consensus gene set was a clear improvement on these individual annotations, with 13,889 genes, 99.1% complete BUSCOs and 86.64% gene loci shared with the reference (Figure 3). Thus, it had a considerably lower percentage of FNs (10.59%) than its RNAseq and homology source annotations, and fewer FPs (2.76%) than all its individual parts. It also had the closest number of predicted genes per locus to the reference annotation number (1.448 OMAnnotator genes per locus:1.458 reference genes per locus; Table 1), underlining its high similarity to the reference annotation.

To compare OMAnnotator with state-of-the-art automated methods, we used two other annotation methods to annotate the *D. melanogaster* genome assembly, with different combinations of source annotations from the earlier testing of OMAnnotator supplied as extrinsic evidence depending on the method’s requirements. The first method, EVM, combines predictions from different sources using weights based on the abundance and source of each prediction. Using the AUGUSTUS, RNAseq and homology annotations as inputs resulted in a consensus annotation with 13,355 genes and 83.26% shared gene loci (Figure 3, Table 1). While these scores indicate a high similarity to the reference annotation, EVM retained fewer gene predictions than OMAnnotator, as indicated by its lower BUSCO Completeness (94.3%, Figure 3) and lower number of average genes per loci (1.251) than the reference (1.314). The EVM annotation also includes significantly more FPs (12.5%, Table 1).

We then ran BRAKER2, which trains the gene finders GeneMark and AUGUSTUS on the target species’ genome assembly, with the option to provide extrinsic homology or RNAseq evidence as hints for training. Using the RNAseq annotation as input produced an annotation with 16365 genes, 96.9% complete BUSCOs and 79.10% shared gene loci (Figure 3, Table 1). Whilst the BUSCO completeness score is high, the gene count suggests overprediction, which is supported by the increase in duplicate BUCSOs (7.8%) and FPs (15.32%). Accordingly, ParsEval analysis shows the average number of BRAKER2 genes per locus (1.425) was higher than the reference number (1.151) (Table 1). We also ran BRAKER2 software with the homology annotation added, to test if this increased the predictive power of the pipeline compared with using RNAseq data alone. However, the output from this run had too many genes (21,816, Figure S2A and B). Running BRAKER2 with the homology annotation alone did not significantly improve specificity and reduced sensitivity (96.5% completeness BUSCOs, Figure S2A). We therefore focus on the annotation produced by incorporating the RNAseq annotation henceforth. Notably, while OMAnnotator had better results in terms of gene content, the percentage of CDS structures matching the reference was slightly higher in the BRAKER2 annotation (81.5%) than for OMAnnotator (79.3%), indicating BRAKER2’s proficiency at modelling gene and exon borders.

### 3.2. OMAnnotator re-annotates three genomes

To illustrate the practical usefulness of OMAnnotator, we re-annotated another three genomes for which source annotations were available: *Siraitia grosvenorii* (Monk Fruit) (Xia *et al*. 2018), *Phyllostomus discolor* (Pale Spear-Nosed Bat) (Jebb *et al*. 2020) and *Olea europaea* (European Olive Tree) (Cruz *et al*. 2016). All of these genomes were originally annotated using EVM with a variety of source annotation numbers and types as input. As there is no gold standard reference annotation for these non-model organisms, EVM and OMAnnotator BUSCO scores are compared with the genome assembly scores (Figure 4).

**Figure 4:**
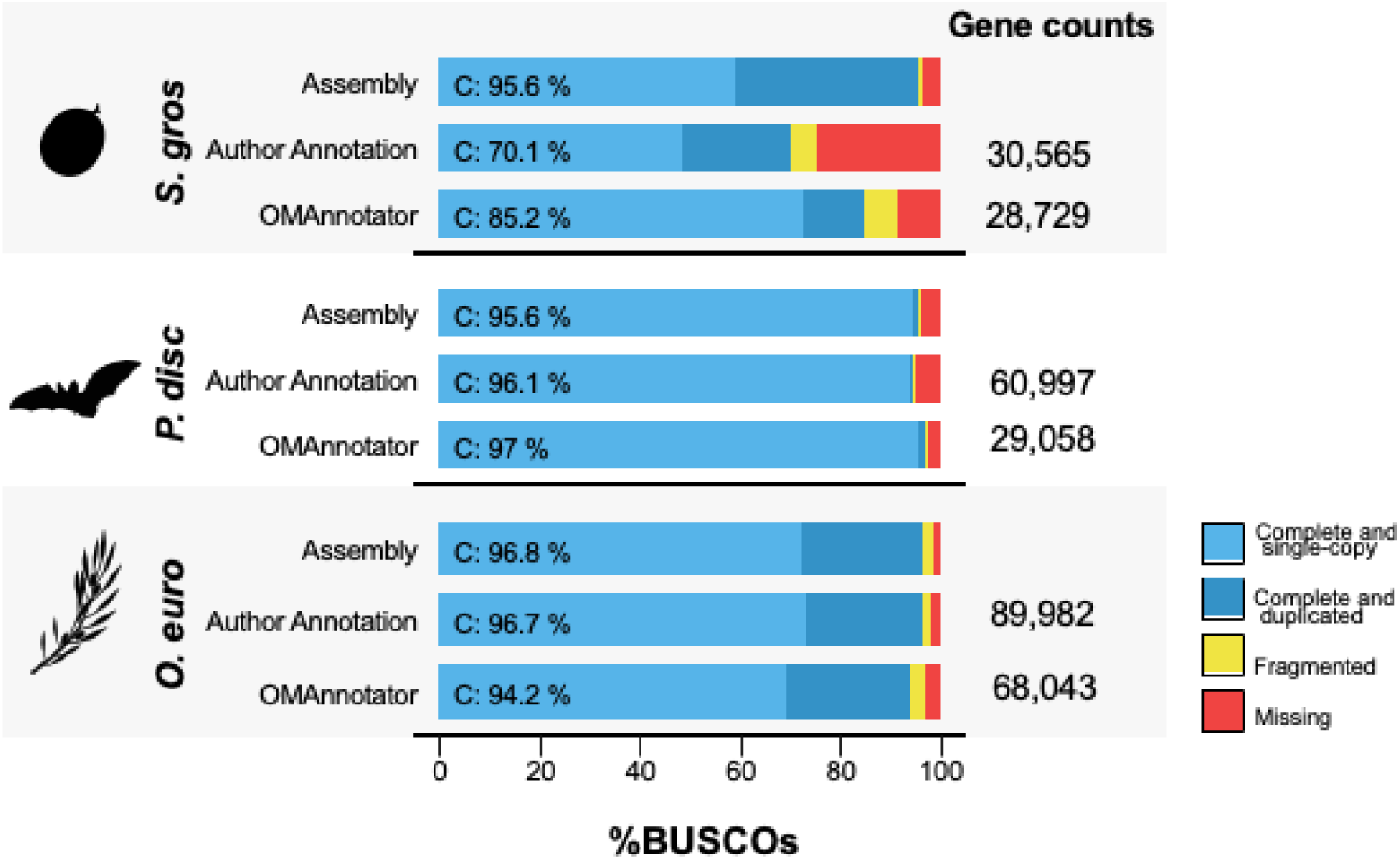
BUSCO results from using OMAnnotator to re-annotate three genomes: Olea europea (top row), Phyllostomus discolour (middle row) and Siratitia grosvenorii (bottom row). The OMAnnotator gene set BUSCO scores are compared to the authors’ annotation set and the genome assembly. Gene counts of the number of genes predicted by each annotation method are shown on the right.

For *S. grosvenorii,* compared to the author’s annotation set of 30,565 genes ((Xia *et al*. 2018)), OMAnnotator predicted 28,729 genes, closer to the mean number of 29,157 genes reported across other *Cucurbitaceae* species (Ma *et al*. 2022). Furthermore the OMAnnotator consensus had fewer missing BUSCO genes, with 97.0% complete BUSCOs – which was much closer to the genome assembly BUSCO completeness score (95.6%) than the EVM consensus (70.1%) (Figure 4).

The *P. discolor* re-annotation followed the same trend: the OMAnnotator consensus had more complete BUSCOs (97.0%) than the author provided EVM consensus (96.1%) and genome assembly (95.6%) (Figure 4). Interestingly, the BUSCO completeness score for the assembly is lower than it is for the annotations. This is a limitation of using the assembly as a benchmark – BUSCO’s genome mode maps BUSCO reference proteins to an assembly, an approach which may not be as sensitive as gene prediction methods. Jebb *et al*. (2020) processed the *P. discolor* EVM annotation further to produce a final annotation with over 99.5% complete BUSCOs and 20,953 genes, which is more in line with the rest of the clade (∼20,000 genes). Based on these numbers, it is likely OMAnnotator retained false genes induced by the nature of the input data (107,272 and 51,137 gene predictions in 2 of their 4 source annotations). Nevertheless, it was less prone to including false positives than EVM.

For *O. europaea*, we were provided with multiple and redundant annotations from the same type of evidence (gene prediction with and without RNA/protein hints) that were used as source annotations by the authors (Cruz *et al*. 2016). Thus, we adapted our species tree input so annotations of the same type were grouped together (see Figure S3). The OMAnnotator annotation had 68,043 genes and 94.2% complete BUSCOs, including 24.7% duplicated. Only the authors’ already manually curated annotation was available for comparison, which was more complete with 89,982 genes and 96.7% complete BUSCOs – closer to the assembly score of 96.8% (Figure 4). Its large number of duplicates is similar to OMAnnotator’s (24.7%) and in line with the assembly (24.5%). Cruz et al. (2016) suggest this large number of duplicates compared to is due to extensive gene duplication in the lamiales lineage leading to the olive tree.

## 4. Discussion

We introduce OMAnnotator, an approach that repurposes OMA Standalone orthology inference software to create a consensus annotation, informed by both individual independent annotations and the gene content of other species. We validated the tool on the *D. melanogaster* genome assembly, demonstrating that the consensus annotation largely outperforms any single input annotation, as well as those made by two other established consensus annotation software run on the same inputs. Finally, we tested the flexibility and robustness of OMAnnotator by using it to re-annotate three genomes that were previously annotated by EVM using varying numbers and types of source annotations.

The quality assessment results for annotations produced by individual methods during proof of principle testing were as anticipated, based on the characteristics of each approach. AUGUSTUS predicts likely genes using a Hidden Markov Model trained with RNAseq and homology data, making it sensitive but prone to false positives. Here, AUGUSTUS was run with the *D. melanogaster* assembly and fly gene model parameters, i.e., a high-quality assembly and a model on which it has been extensively trained — its optimal use case. Nevertheless, it had the most unique predictions compared to the reference. Additionally, even with its high sensitivity, the number of missing genes was significantly higher than the OMAnnotator consensus. RNAseq data contains what is expressed in the tissues at the time of extraction, meaning genes expressed at different developmental stages can be missed, resulting in a set with more fragmented and missing genes relative to other methods. Finally, the success of homology annotation depends on the degree of relatedness between the target and the reference species and the quality of the reference proteome. In this study, we chose the *A. gambiae* proteome as our reference to mimic the situation in which the novel genome is from a species with no closely related model organism, thereby producing an annotation with many missing genes.

The OMAnnotator consensus improves on these individual methods, filtering false gene predictions and combining true genes into a more complete consensus annotation closer to the reference than any individual method. One caveat to the BUSCO result is that OMAnnotator uses orthology data from other species to find evidence for including predicted genes in the consensus. It is therefore expected to favour predictions with highly conserved orthologs, and thus, it is expected to obtain relatively high BUSCO completeness. Nevertheless, as both the gene count and ParsEval analysis support the BUSCO result, we can be confident that the favourable BUSCO score is not solely due to this bias.

OMAnnotator also performed well compared to two other state-of-the-art pipelines: EVM and BRAKER2, which were chosen based on their popularity and availability of documentation (Haas *et al*. 2008; Hoff *et al*. 2019). EVM was less prone to overprediction than BRAKER2, perhaps due to its different use case: combining gene models from the user-inputted annotations, dropping spurious predictions, rather than using them as extrinsic evidence for training gene finder software. The most recent release of BRAKER3, unavailable at the time of our study, has the <u>T</u>ranscript <u>SE</u>lector for <u>BRA</u>KER tool integrated into its pipeline, which likely reduces the overprediction rate (Gabriel *et al*. 2021, 2023). Another note is that although OMAnnotator is more accurate for gene content prediction, BRAKER2 appears more accurate in its delimitation of gene intron-exon structure, which is not modelled explicitly by OMAnnotator. Since OMAnnotator can combine predictions from any GFF, it is not exclusive to any specific combination of annotation methods. We therefore propose that OMAnnotator be used to combine annotations from methods that are specially designed to accurately predict exon-intron structures (such as BRAKER2/3) with other source annotations to increase sensitivity without sacrificing accuracy. Overall, OMAnnotator produced the annotation closest to the reference during proof of principle testing, achieving an excellent sensitivity-to-specificity balance as an annotation combiner.

Any new genome annotation software designed to combine gene predictions must be flexible enough to accept multiple types of annotation sets across many species as input and robust enough to still filter FPs. We demonstrate that OMAnnotator possesses these qualities by re-annotating three non-model organism genomes using the authors’ source annotation sets, which were variable in number and type. It consistently predicted a number of genes similar to expectations, while maintaining BUSCO completeness scores in line with the assembly completeness. Moreover, OMAnnotator was improved upon the EVM outputs for *S. grosvenorii* and *P. discolor*, underscoring the usefulness of leveraging OMA’s orthology data. In short, OMAnnotator keeps its specificity-to-sensitivity balance even with numerous source annotations for non-model species.

Although OMAnnotator produced the highest quality first annotations, the final annotations for and *O. europaea* had better results overall. The *P. discolor* final annotation reported by Jebb et al. (2020) had undergone additional rounds of PASA filtering, reducing the predicted gene count in line with expectations based on the rest of the clade and improving BUSCO results. The *O. europaea* final annotation produced by Cruz et al. (2016) had very few missing genes due to author curation, which OMAnnotator was unable to match despite producing a high quality gene repertoire according to BUSCO. Thus, while OMAnnotator produces highly complete and accurate first annotations by utilising orthology evidence, downstream curation remains an important part of refining an annotation.

During our benchmarking of OMAnnotator performance, we strove to acquire unprocessed annotations to allow fair comparison between EVM and OMAnnotator annotations. Though we achieved this for two out of three species, only the authors’ final *O. europaea* annotation was available for comparison – we cannot know if OMAnnotator produced a higher quality first annotation. Several of the re-annotation candidates we identified did not have all annotations from intermediate steps publically available (or even available on request), limiting the scope of our analysis. This highlights the importance of improving reproducibility in bioinformatics pipelines, by releasing not only final data but also intermediary steps.

To conclude, by repurposing the OMA algorithm to use evolutionary data for building consensus annotations, we have created a potent new tool in the genome annotation toolbox — OMAnnotator. Its development is timely due to the continued rise of genome sequencing (Hotaling, Kelley and Frandsen 2021; Marks *et al*. 2021) and the widely recognised challenge of eukaryotic genome annotation (Salzberg 2019). As OMAnnotator makes use of existing gene repertoires in evolutionary databases, its accuracy will increase as more high quality gene annotations are made available. We expect it to become a new step in annotation pipelines, where it will benefit from ongoing efforts to develop and improve gene finder methods (Scalzitti *et al*. 2020), whose annotations it can use as input datasets.

## Data Availability

OMAnnotator is available on GitHub (https://github.com/DessimozLab/OMAnnotation). The OMAnnotator and OMA standalone versions used for this study are available on Zenodo, along with all analyses (https://doi.org/10.5281/zenodo.14252311).

## Supplementary information

Supplementary information is available in the accompanying Supplementary Material document.

## Author contributions

YN and CD conceptualised and supervised the study. YN wrote the software. SB and YN carried out the investigation and analysis. SB and YN wrote the manuscript. SB, YN and CD edited, reviewed and approved the final version of the manuscript.

## Acknowledgements

We thank Adrian Altenhoff for his technical assistance using OMA Standalone tools and Clemént Train for his help with using pyHAM. We are grateful for all the responses we received from the authors we contacted regarding their genome annotations. Particular thanks go to the authors whose annotation data we used for this study: Michael Hiller, who shared the BAT1K annotation dataset, Xue Han, who shared the *Siraitia grosvenorii* annotation set, and Tyler Alioto, who shared the *Olea europaea* annotation dataset.

