## Supplementary for "OMAnnotator: a novel approach to building an annotated consensus genome sequence"

### Supplementary Material

**Figures**

Figure S1: Species tree used for OMAnnotator proof of principle testing

Figure S2: BUSCO results from BRAKER2 *D. melanogaster* annotations with RNAseq only (BRAKER2 RNAseq), homology only (BRAKER2 pep) and RNAseq plus protein evidence (BRAKER2 RNA plus pep).

Figures S3-S6: Species trees used for OMAnnotator re-annotations

**Tables**

Table S1: Lists of related species used for OMAnnotator annotations

#### Supplementary Figures

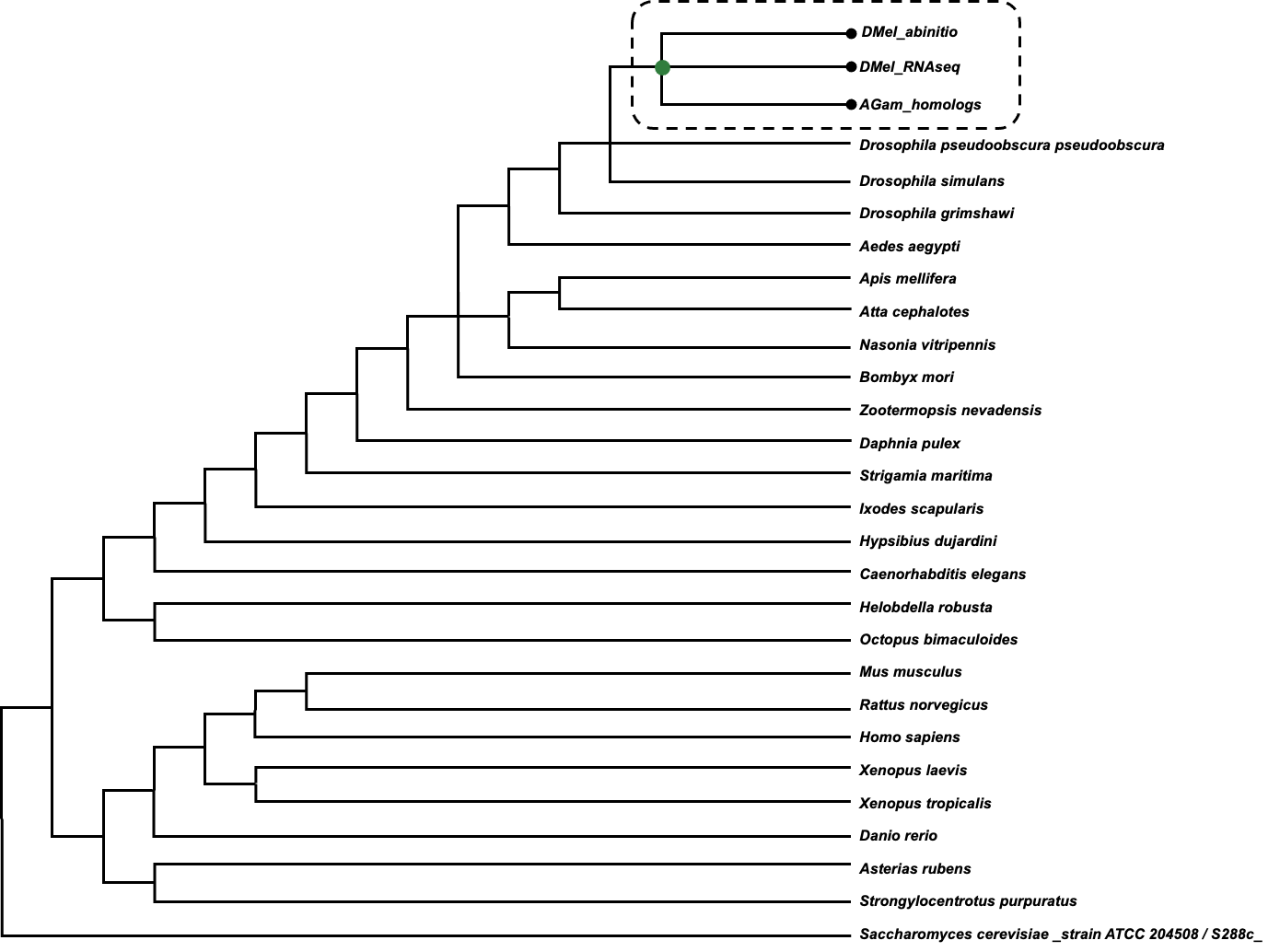

*Figure S1: Proof of principle D. melanogaster OMAnnotator species tree. Source annotations are highlighted by the dashed box and black circles. The green circle shows the node at which the consensus annotation is constructed.*

***A)***
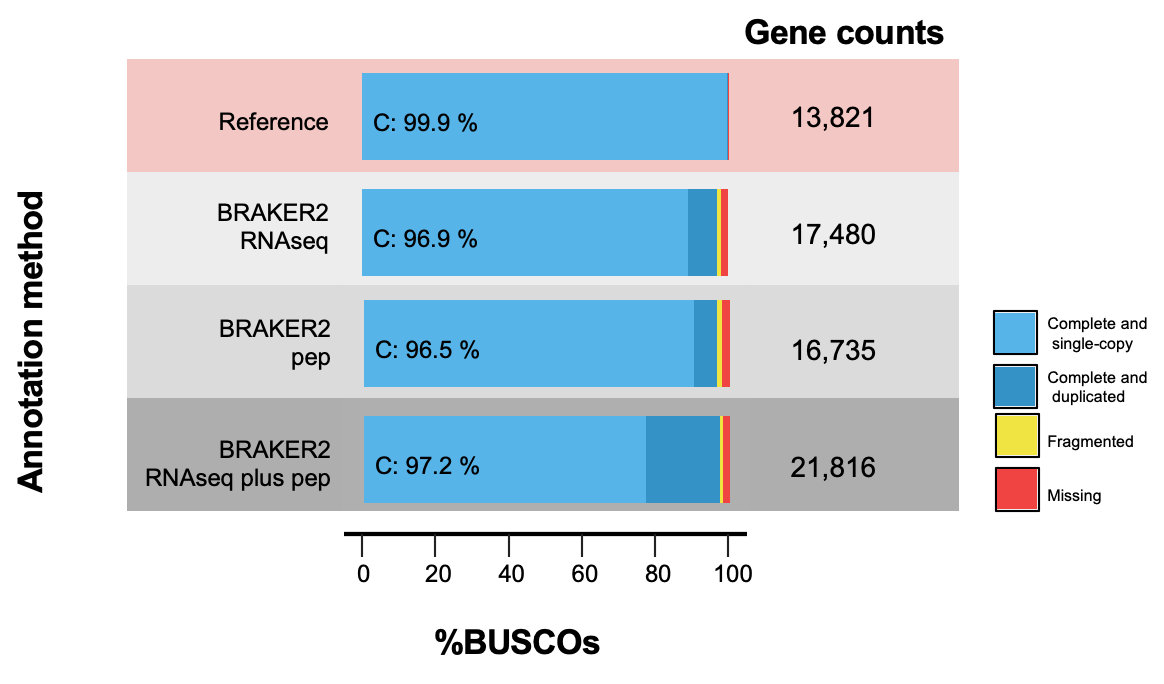

***B)***

| ***ParsEval comparison level*** | ***Comparison*** | ***BRAKER2***  ***RNAseq*** | ***BRAKER2***  ***pep*** | ***BRAKER2***  ***RNA plus pep*** |
| --- | --- | --- | --- | --- |
| *Genes* | *Total number* | *16,365* | *16,282* | *21,816* |
| *Gene Loci* | *Shared (%)* | *79.10%* | *79.01%* | *80.73%* |
|  | *Unique to reference (FN)* | *4.03%* | *5.45%* | *3.95%* |
|  | *Unique to prediction (FP)* | *16.87%* | *15.53%* | *15.32%* |
|  | *Genes per locus reference* | *1.151* | *1.156* | *1.179* |
|  | *Genes per locus prediction* | *1.425* | *1.370* | *1.832* |
| *CDS structure* | *Prediction CDS matches to reference* | *81.5%* | *78.9%* | *81.2%* |

*Figure S2: Quality assessment results for all three BRAKER2 annotations of the D. melanogaster genome assembly.*

*Annotations were performed using RNAseq data only (BRAKER2 RNAseq), homology data only (BRAKER2 pep), and RNAseq and protein data (BRAKER2 RNA plus pep). A) BUSCO results and gene counts for each BRAKER2 annotation are shown in grey shades. The reference annotation BUSCO results and gene counts are highlighted in red at the top. B) ParsEval results for each BRAKER2 annotation are shown. BRAKER2 RNAseq results are reported in the main manuscript (as BRAKER2), as they are better than BRAKER2 with RNAseq and protein data, and RNAseq data is more likely to be available as an OMAnnotator source annotation than protein data only.*

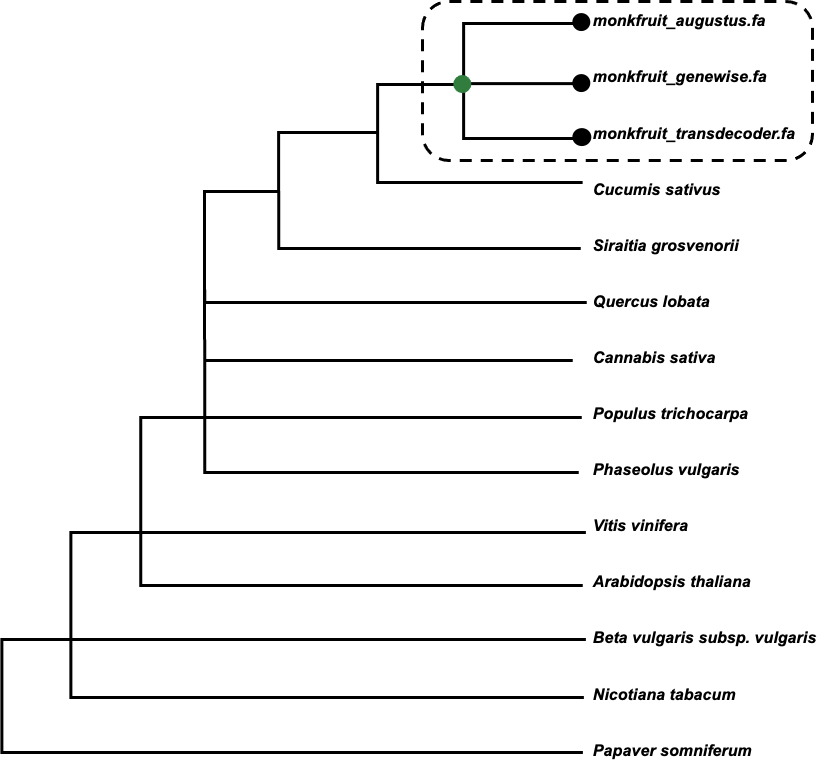

*Figure S4: Siraitia grosvenorii OMAnnotator species tree. Source annotations are highlighted by the dashed box and black circles. The green circle shows the node at which the consensus annotation is constructed.*

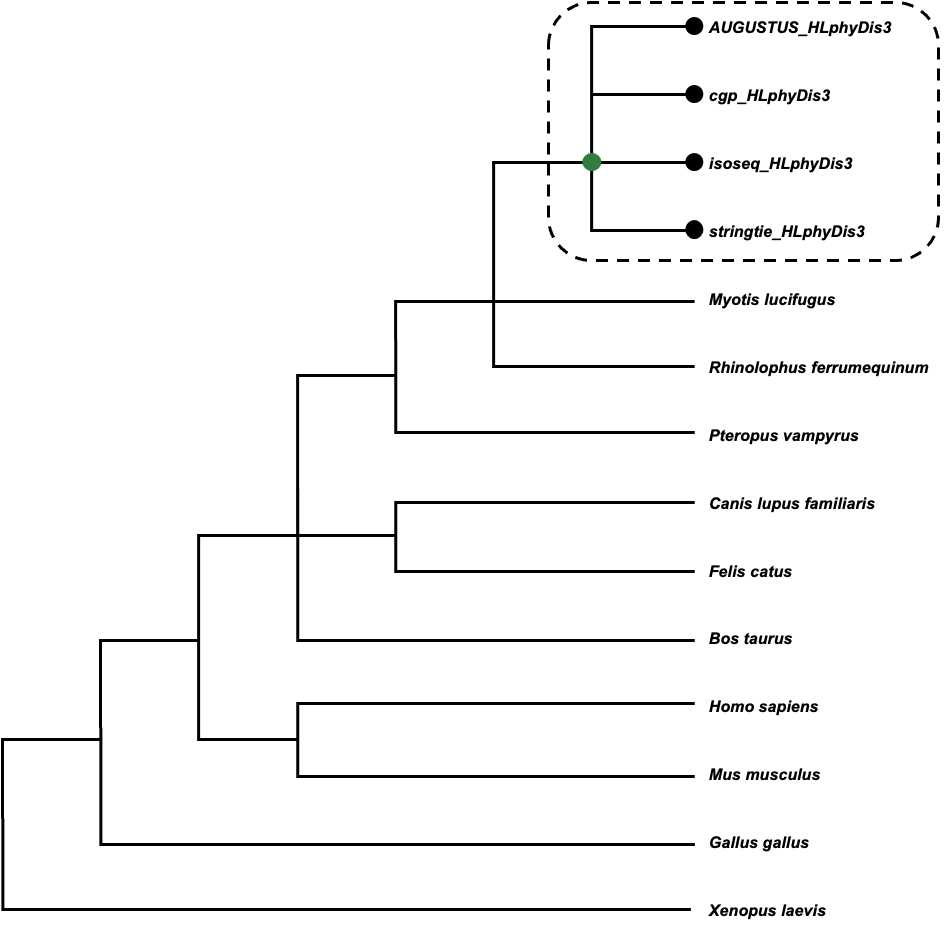

*Figure S5: Phyllostomus discolor OMAnnotator species tree. Source annotations are highlighted by the dashed box and black circles. The green circle shows the node at which the consensus annotation is constructed.*

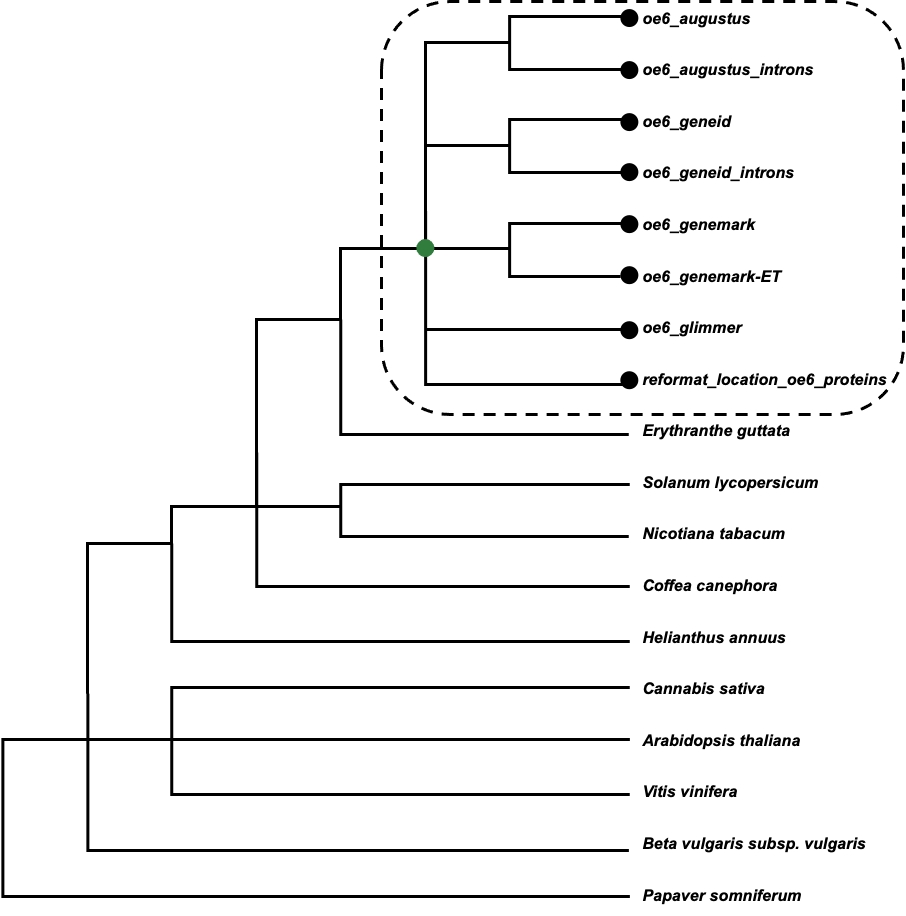
*Figure S3: Olea europaea OMAnnotator species tree. Source annotations are highlighted by the dashed box and black circles. The green circle shows the node at which the consensus annotation is constructed.*

#### Supplementary Tables

*Table S1: Lists of species selected to form the precomputed set of orthology relationships for each OMAnnotator annotation. These data were downloaded from the OMA Browser (*[*https://omabrowser.org/oma/export/*](https://omabrowser.org/oma/export/)*) with the OMA standalone software, forming each species' OMA folder.*

| ***Species included in precomputed orthology relationships dataset*** | | | | |
| --- | --- | --- | --- | --- |
| ***Species number*** | ***Drosophila melanogaster*** | ***Siraitia grosvenorii*** | ***Phyllostomus discolor*** | ***Olea europaea*** |
| *1* | *Caenorhabditis elegans* | *Papaver somniferum* | *Xenopus laevis* | *Papaver somniferum* |
| *2* | *Ixodes scapularis* | *Beta vulgaris subsp. vulgaris* | *Gallus gallus* | *Beta vulgaris subsp. vulgaris* |
| *3* | *Strigamia maritima* | *Nicotiana tabacum* | *Myotis lucifugus* | *Helianthus annuus* |
| *4* | *Daphnia pulex* | *Arabidopsis thaliana* | *Rhinolophus ferrumequinum* | *Solanum lycopersicum* |
| *5* | *Bombyx mori* | *Vitis vinifera* | *Pteropus vampyrus* | *Nicotiana tabacum* |
| *6* | *Aedes aegypti* | *Cannabis sativa* | *Bos taurus* | *Erythranthe guttata* |
| *7* | *Drosophila grimshawi* | *Cucumis sativus* | *Canis lupus familiaris* | *Coffea canephora* |
| *8* | *Drosophila simulans* | *Populus trichocarpa* | *Felis catus* | *Cannabis sativa* |
| *9* | *Drosophila pseudoobscura pseudoobscura* | *Phaseolus vulgaris* | *Homo sapiens* | *Arabidopsis thaliana* |
| *10* | *Nasonia vitripennis* | *Quercus lobata* | *Mus musculus* | *Vitis vinifera* |
| *11* | *Apis mellifera* |  |  |  |
| *12* | *Atta cephalotes* |  |  |  |
| *13* | *Zootermopsis nevadensis* |  |  |  |
| *14* | *Hypsibius dujardini* |  |  |  |
| *16* | *Helobdella robusta* |  |  |  |
| *17* | *Octopus bimaculoides* |  |  |  |
| *18* | *Danio rerio* |  |  |  |
| *19* | *Xenopus laevis* |  |  |  |
| *20* | *Xenopus tropicalis* |  |  |  |
| *21* | *Homo sapiens* |  |  |  |
| *22* | *Mus musculus* |  |  |  |
| *23* | *Rattus norvegicus* |  |  |  |
| *24* | *Asterias rubens* |  |  |  |
| *25* | *Strongylocentrotus purpuratus* |  |  |  |
| *26* | *Saccharomyces cerevisiae*  *_strain ATCC 204508 / S288c_* | |  |  |
